# Heparan sulfate promotes autoactivation of pro-Cathepsin K by destabilizing the propeptide-catalytic domain interaction

**DOI:** 10.64898/2026.01.18.700217

**Authors:** Huanmeng Hao, Xiaoxiao Zhang, Jian Liu, Ding Xu

**Affiliations:** Department of Orthopedics, School of Medicine, Emory University; Department of Biochemistry, School of Medicine, Emory University; Department of Oral Biology, School of Dental Medicine, University at Buffalo, the State University of New York.; Division of Chemical Biology and Medicinal Chemistry, Eshelman School of Pharmacy, University of North Carolina

**Author notes:** To whom correspondence and proofs should be sent: Department of Orthopaedics, School of Medicine, Emory University, Atlanta, GA30329.

**Keywords:** Protease, glycosaminoglyan, cathepsin, osteoclasts, lysosome

## Abstract

Pro-cathepsin-K (pro-CtsK) is the zymogen of cathepsin-K (CtsK), a collagenase that is essential for bone resorption. pro-CtsK is known to bind heparan sulfate (HS), but the biological significance of the interaction remains unclear. Here we report that HS accelerates the autoprocessing of pro-CtsK in a manner dependent on both sulfation pattern and oligosaccharide length. We discovered a previously unknown electrostatic interaction between the propeptide and the catalytic domain, which stabilizes the conformation of the propeptide and prevents it from intermolecular proteolytic activation. HS accelerates autoprocessing of pro-CtsK by disrupting this critical electrostatic interaction. Mechanistically, HS competes with two glutamic acids in the propeptide for binding to three basic residues on the catalytic domain, thereby substantially alters the conformation of the propeptide and making it more labile for autoprocessing. We further discovered that HS is highly enriched in secretory lysosomes of osteoclasts and might be directly involved in autoactivation of CtsK.

## Introduction

Cathepsin K (CtsK) is a member of the cysteine proteases family highly expressed by mature osteoclasts and is essential for bone resorption (1). Among all known cathepsins, CtsK is unique in that it possesses exceptionally potent collagenase activity. It efficiently degrades type I collagen, the primary component of the organic bone matrix (2). The physiological importance of CtsK in bone homeostasis has been evidenced in pycnodysostosis, a rare autosomal recessive human disease caused by mutations in *ctsk* (*3–5*). Patients exhibit osteosclerosis, short stature, and skeletal fragility, demonstrating tightly regulated CtsK activity is required for maintaining skeletal health(6).

Like other cysteine proteases, CtsK is synthesized as an inactive precursor (pro-CtsK) that undergoes proteolytic processing to generate mature and enzymatically active enzyme (7). Pro-CtsK activation favors acidic conditions (∼pH 4) and requires removal of the N-terminal pro-domain through autoactivation or by other proteases. Acidic pH induces a conformational change in pro-CtsK that unmasks the active site and renders the zymogen susceptible to autoproteolytic processing (8). Because zymogen activation is the first and rate-limiting step in controlling protease activity, pro-CtsK maturation is expected to be tightly regulated.

Glycosaminoglycans (GAGs) are negatively charged linear polysaccharides capable of binding hundreds of proteins mainly through electrostatic interactions with basic residues (9). Many cysteine proteases bind GAGs and their activities can be regulated by different types of GAGs (10). It has been shown that in the presence of GAGs, the autoactivation process of several cathepsins, including cathepsin B, L, and K, are greatly accelerated (11–14). For both cathepsin B and L, similar mechanisms were proposed where direct binding of GAGs to basic residues on the prodomain induces a conformational change of the prodomain, converting the propeptide into a better substrate to allow more efficient digestion through intermolecular proteolytic cleavage(13, 14). Similarly, it was shown that binding of Chondroitin-4-sulfate (C4S) to proCtsK induces a conformational change, will likely allow more efficient autoactivation(11). However, for all these studies structural insights remain lacking regarding how GAGs induce such conformational changes and why the altered conformation allows more efficient digestion.

In this study, we discovered that heparan sulfate (HS) greatly accelerates pro-CtsK autoactivation, in a manner dependent on both oligosaccharide length and sulfation level. Through mutagenesis studies, we identified several electrostatic interactions between the propeptide and the catalytical domain that play essential roles in protecting pro-CtsK from autoactivation. Interestingly, the arginine residues involved in these electrostatic interactions in the catalytic domain also makes essential contribution to HS binding. These findings support a model in which HS directly competes with the two glutamic acid residues in the propeptide for binding to the arginine residues in the catalytic domain, thereby destabilizing the conformation of the propeptide and facilitate autoactivation. Finally, immunofluorescence analysis of osteoclasts in bone sections revealed substantial co-localization of HS and CtsK within secretary lysosomes, indicating a potential role for HS in promoting pro-CtsK maturation in vivo. Collectively, these findings delineate the structural details by which HS modulates CtsK activation and provide new biochemical insights of autoactivation of cysteine cathepsins.

## RESULTS

### Heparin markedly accelerates autoprocessing of pro-mCtsK

To determine whether heparin promotes the autoprocessing of pro-mCtsK, we incubated pro-mCtsK in autoactivation buffer (pH 4) at room temperature for up to 6 hrs in the presence or absence of a 1:1 molar ratio of heparin. Heparin dramatically increased the rate of pro-mCtsK autoprocessing, enhancing conversion to mature form by approximately 6-fold relative to the no-heparin control (Fig. 1A). Consistently, peptidase activity measurement using a peptide substrate confirms the autoprocessing rate was increased by more than 6-fold (Fig. 1B).

**Figure 1.**
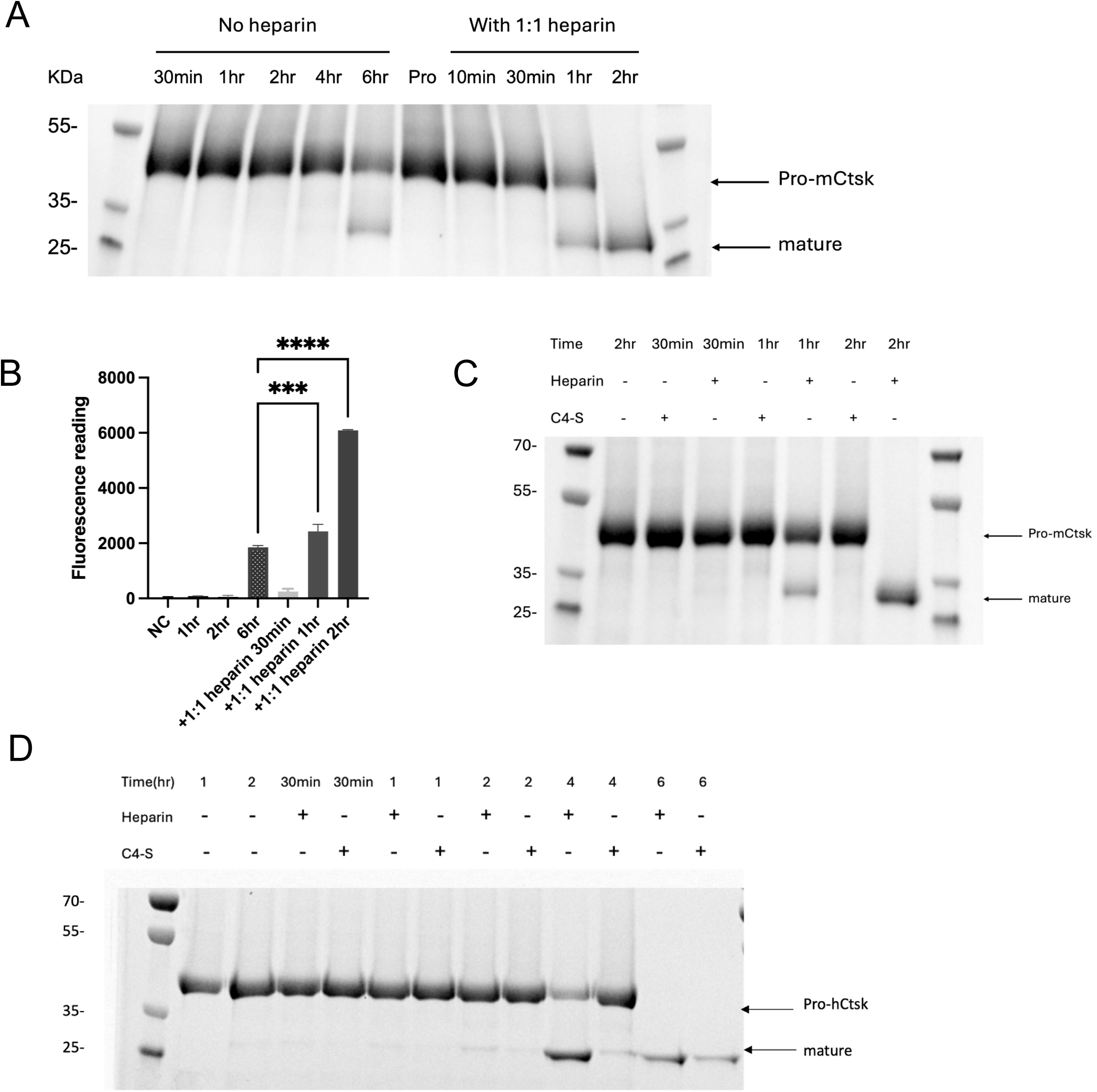
Heparan sulfate promotes pro-Cathepsin K autoactivation at acidic pH. (A) Pro-mCtsK was incubated with or without heparin (1:1 molar ratio) at pH 4. The reaction was performed at 22 °C and monitored by commassie blue staining. (B) CtsK activity in the autoactivation reactions (same conditions as in panel A) was measured by fluorescent peptide substrate Z-Leu-Arg-AMC. (C) Autoprocessing of pro-mCtsk was monitored in the presence or absence of heparin and C4-S (22 °C, pH 4). (D) Autoprocessing of human pro-cathepsin K (pro-hCtsK) was monitored in the presence or absence of heparin and C4-S (37 °C, pH5). Data representative of at least three similar experiments.

Because C4S has previously been reported to promote autoactivation of human CtsK (11), we directly compared C4S with heparin under identical conditions. Heparin accelerated pro-mCtsK processing significantly more efficiently than C4S (Fig. 1C). We further confirmed that C4S was indeed able to enhance human procathepsin K (pro-hCtsK) autoprocessing at 37 °C, pH 5 as reported (11), but again this promoting effect lags behind heparin under the same condition (Fig. 1D).

### Sulfation and length requirements for HS-mediated activation of mCtsK

Previously, we have shown that the minimum length of HS oligosaccharide required for forming a stable complex with CtsK is dodecasaccharide (12mer) (15). To define the structural requirements for HS-mediated pro-mCtsK autoactivation, we examined a panel of HS-12mer oligosaccharides differing in sulfation levels (NS2S, NS2S6S, and NS2S3S6S). The minimally sulfated 12mer-NS2S only moderately accelerates the autoactivation of mCtsK relative to control, whereas 12mer-NS2S6S markedly increased the rate of pro-mCtsK processing. 12mer-NS2S3S6S contains only one additional sulfate group at the 3-O-position of one of the glucosamine residues, but it displayed clear enhancement of autoprocessing at 1 hour compared to 12mer-NS2S6S. (Fig. 2A). These findings demonstrate a clear sulfation-dependent manner in HS-mediated activation and highlights the sensitivity of HS-mediated CtsK autoprocessing to HS sulfation patterns.

**Figure 2.**
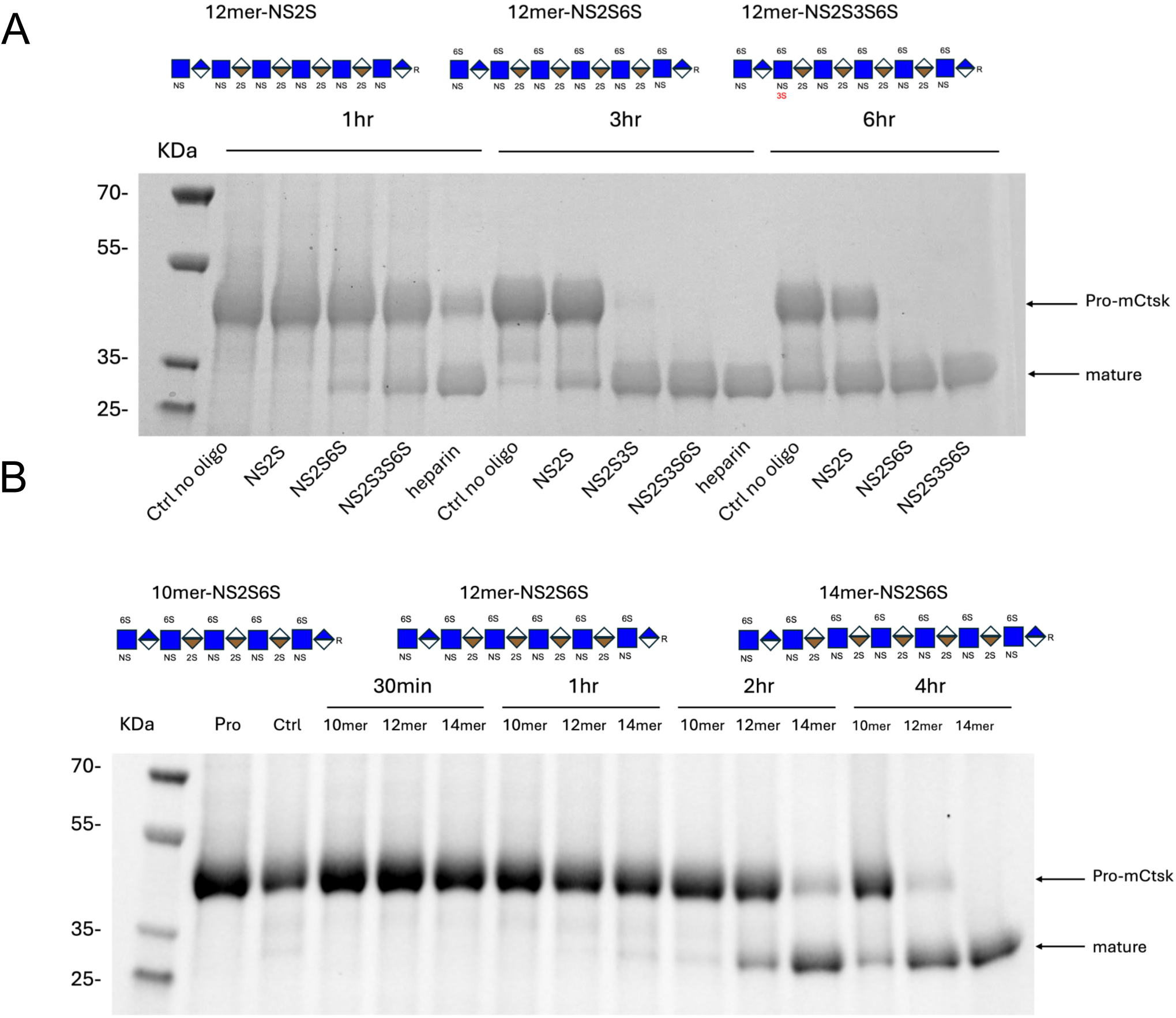
Heparan sulfate oligosaccharides promotes pro-CtsK autoactivation in a sulfation-dependent and length dependent manner. (A) Autoprocessing of pro-mCtsk in the presence of three 12mer HS oligosaccharides with different sulfation levels at 1, 3 and 6 hours. (B) Autoprocessing of the pro-mCtsk in the presence of 10mer, 12mer, and 14mer HS oligosaccharides from 30 minutes to 4 hours. In all experiments, pro-mCtsk was incubated with oligosaccharides at 1:1 molar ratio at pH 4, 22 °C. Image representative of at least three similar experiments.

We next investigated the contribution of HS length by comparing 10-, 12-, and 14-mer oligosaccharides. As shown in Fig. 2B, it is apparent that the autoactivation process facilitated by 14-mer is the fastest, almost completing process within 2hrs. After 4hrs, both 12-mer and 14-mer could fully convert pro-mCtsK to its mature form, whereas 10-mer showed minimum promotion effect. These results indicate that longer HS oligosaccharides are more effective in activating mCtsK, and even slight changes in the length has a big impact on the rate of autoprocessing.

### HS-binding likely causes a steric clash between HS and propeptide

Having defined the structural requirements for HS-mediated activation, we next sought to understand how HS binding may alter pro-mCtsK conformation. Recently, we have solved the co-crystal structure of mature mCtsK in complex with 12mer-NS2S6S (PDB: 8V58) and identified the residues that directly interact with HS (Fig. 3A) (15). Overlay of this structure with the structure of pro-hCtsK (PDB:1BY8) revealed a pronounced steric clash between the propeptide loop and the bound HS oligosaccharide (Fig. 3B), suggesting that HS binding would require, or induce, a conformational change of pro-mCtsK that involves a large swing of the loop away from the bound HS. This structural insight supports the hypothesis that HS-binding induces/selects a more labile conformation of the propeptide loop, thereby making it a better substrate for autoprocessing by intermolecular proteolytic cleavage.

**Figure 3.**
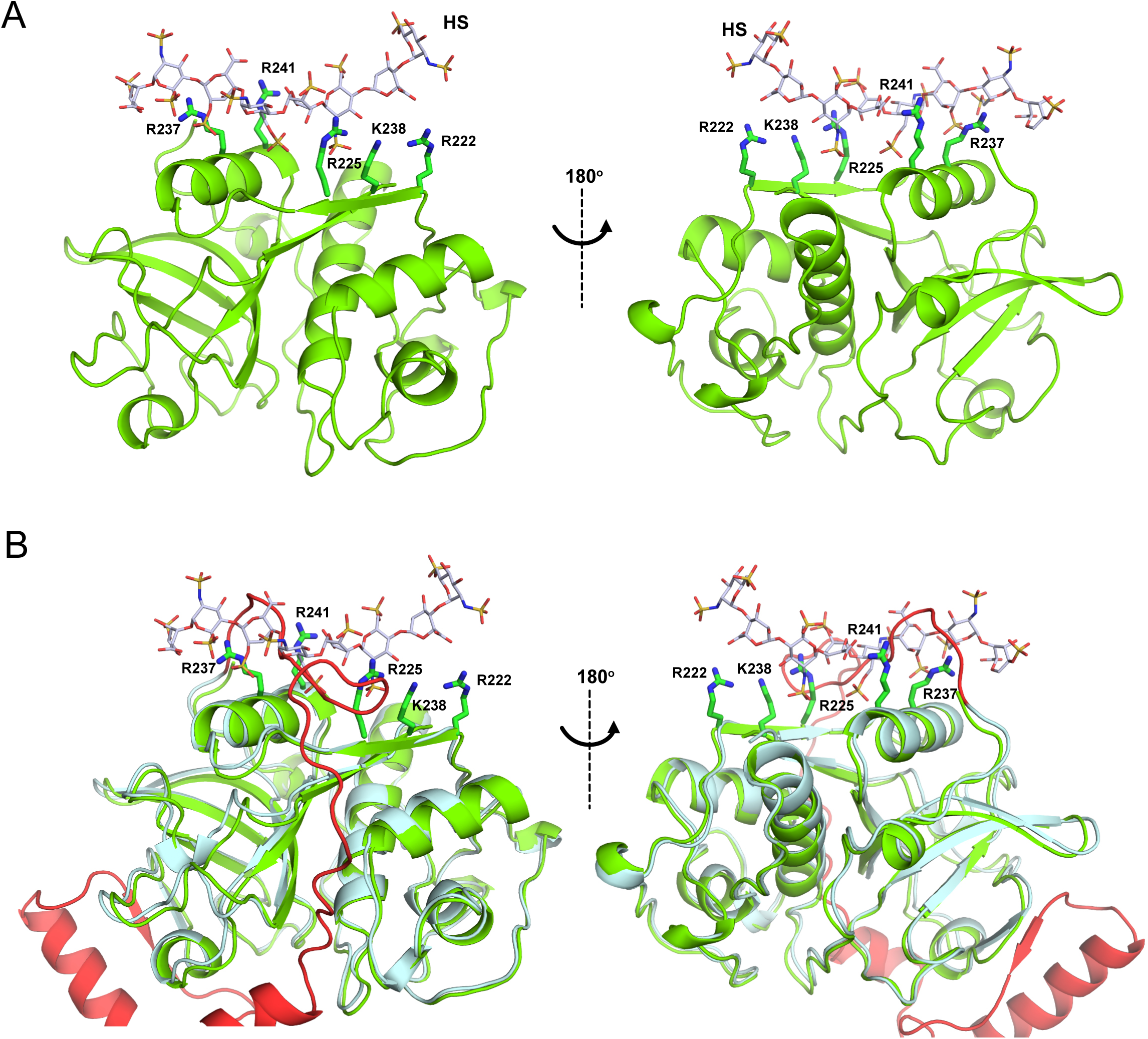
HS-binding likely causes a steric clash between HS and propeptide. (A) Crystal structure of mCtsK-HS complex (PDB: 8V58). Mature mCtsk (green) is shown in cartoon representation. The HS oligosaccharide is shown as sticks (carbon backbone in grey, sulfur in yellow, oxygen in red and nitrogen in blue). Residues making major contributions to HS-binding are shown in sticks. (B) Overlay of the structures of human procathepsin K (pro-hCtsk) and mCtsk-HS complex. The propeptide of pro-hCtsk is shown in red and the catalytic domain is shown in cyan. Steric clash of the propeptide connection loop and the HS oligosacchride is observed above the a-helix containing residues R237 and R241.

### Disruption of electrostatic interactions within the connection loop enhances pro-mCtsK autoactivation

In the pro-hCtsK crystal structure the last stretch of the connection loop between the prodomain and the mature enzyme (T105-A115) sits right above three HS-binding residues (R225, R237 and R241) that make the most prominent contribution of HS-binding. Interestingly, this stretch contains two glutamic acids (E110 and E112) that are located in very close proximity to R237 and R241 (Fig. 4A). Homology alignment found that the negative charges at these two positions are completely conserved among all mammals with some species having an aspartic acid at the 110 position. This finding prompt us to hypothesize that the conformation of the connection loop might be stabilized by electrostatic interactions between E110 and E112 and R225, R237 and R241.

**Figure 4.**
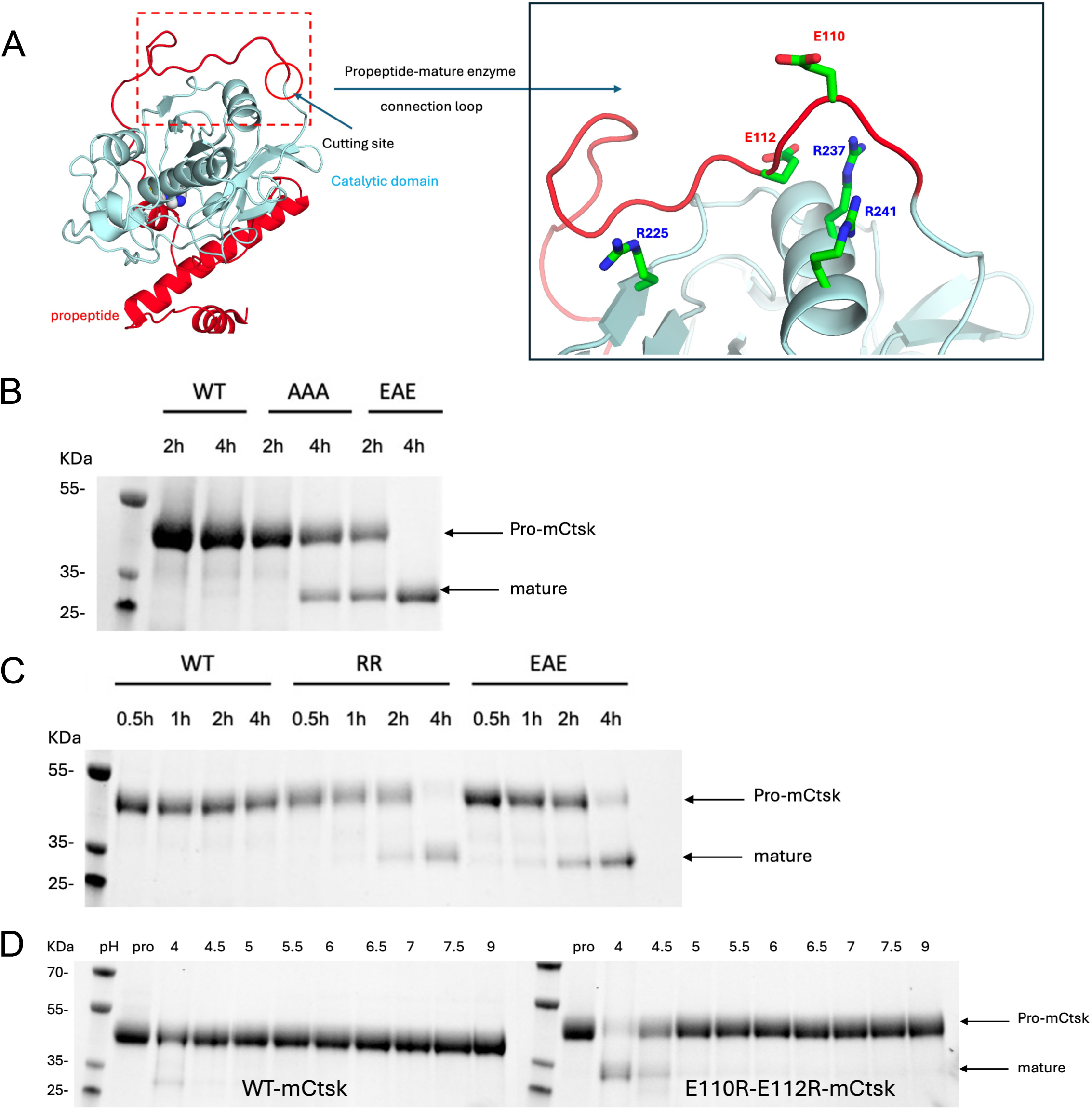
Disruption of electrostatic interactions between the propeptide and the catalytic domain accelerates pro-mCtsK autoactivation. (A) Proposed electrostatic interactions between the propeptide connection loop (red) and the catalytic domain a-helix (cyan) based on the structure of pro-hCtsK (PDB:1BY8). (B) Autoprocessing of pro-mCtsK WT, R225A-R237A-R241A (AAA), and R225E-R237A-R241E (EAE) at 22 C, pH4. (C) Autoprocessing of pro-mCtsK WT, E110R-E112R (RR) and EAE mutants at 22 C, pH4. (D) Autoprocessing of WT pro-mCtsk (left) and E110R/E112R double mutant (right) for 4 hrs at 22C in different pH.

To test our hypothesis, the involvement of these charged residues in stabilizing proCtsK conformation was investigated by mutagenesis. We manipulated the charges in this connection loop to disrupt the proposed electrostatic interactions. Three mutants were generated by site-directed mutagenesis, including E110R-E112R (RR), R225A-R237A-R241A (AAA), and R225E-R237A-R241E (EAE). The idea here is to swap the charges of propepetide or mature domain to weaken the intramolecular interactions or introduce repelling charges. The processing of various mutants was investigated in the absence of HS in acidic buffer at room temperature. As shown in Fig. 4B, both AAA and EAE mutants autoprocessed substantially faster than WT, with the EAE mutant reaches complete autoprocessing by 4 hours. Strikingly, RR mutant and EAE mutants, both bearing opposite charges to the native residues, underwent highly similar rate of acceleration of autoprocessing (Fig. 4C). This result strongly suggests the importance of the electrostatic interaction between E110/E112 and R225/R237/R241 in maintaining the zymogen form. Furthermore, while WT pro-mCtsK showed no detectable processing across pH4.0-9.0 at RT for 4hrs, the E110R-E112R mutant displayed a shift toward activation at pH4.5, further highlighting its destabilized conformation. Overall, our data indicates that disruption of the electrostatic interactions in the connection loop accelerates pro-mCtsK autoactivation and point to this region as the potential target of HS-mediated destabilization.

Having established that the electrostatic interactions between the acidic residues in the connection loop and the basic residues in the catalytic domain play critical role in regulating autoprocessing of pro-CtsK, we wonder whether the same mechanism might also in play in other cathepsins. Interestingly, in both procathepsin B and L, we found similarly positioned acidic residues in the propeptide and basic residues in the catalytic domain (Fig. 5). The presence of these potential electrostatic interactions suggests that procathepsin B and L might also use similar mechanism to regulate their autoactivation process.

**Figure 5.**
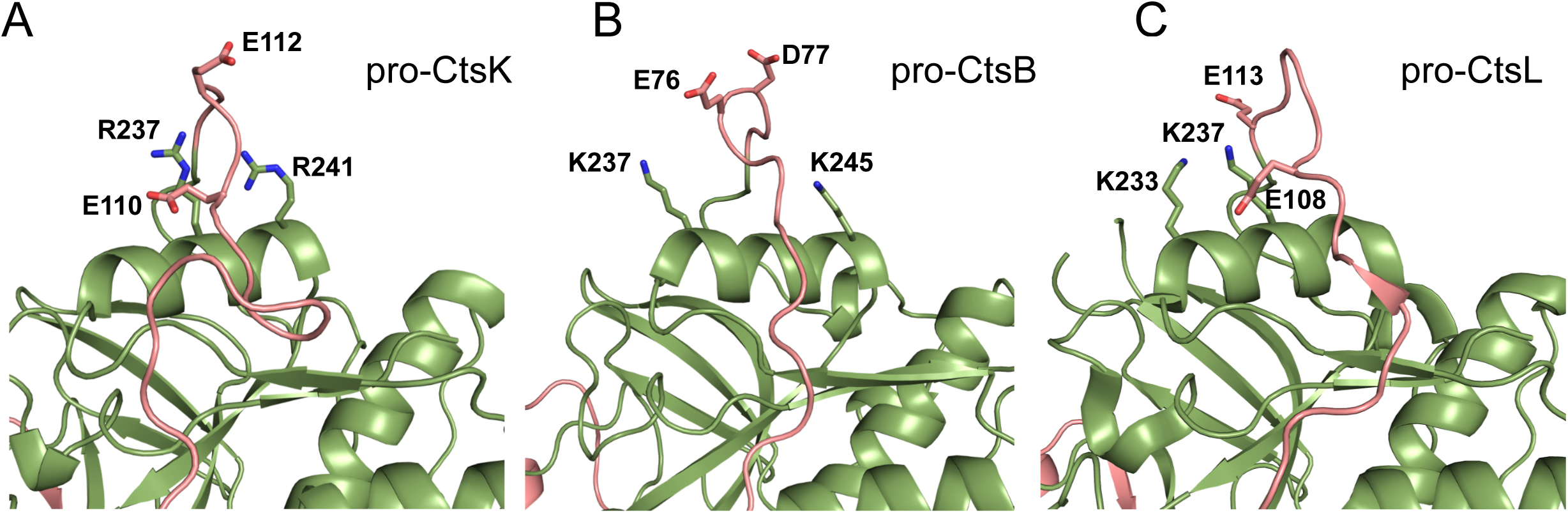
Procathepsin B (pro-CtsB) and procathepsin L (pro-CtsL) also contains similar arrangement of acidic residues in the connection loop and basic residues in the catalytic domain. (A) Cartoon representation of human proCtsk structure (PDB:1BY8). (B) Cartoon representation of human proCtsB structure (PDB:3PBH). (C) Cartoon representation of human proCtsL structure (PDB:6JD8).The propeptide domains for all structures are shown in salmon and the catalytic domains shown in green.

### HS-binding destabilize pro-mCtsK conformation

To further confirm our hypothesis that HS-binding induces/selects a more labile conformation of the propeptide loop by disrupting the electrostatic interactions between E110/E112 and R225/R237/R241, we examined the thermostability of pro-mCtsK in the presence of absence of heparin using nanoscale Differential Scanning Fluorimetry (nanoDSF). We found heparin-binding resulted in a ∼8 °C reduction in Tm value, indicating greatly reduced thermostability of pro-mCtsK (Fig. 6A). Interestingly, E110R/E112R mutant displayed a highly similar 8 °C reduction in Tm value compared to WT pro-mCtsK (Fig. 6B). Combined, these results strongly suggest that HS promotes proCtsK autoactivation through disturbing the electrostatic interactions between E110/E112 and R225/R237/R241, which play a critical role in protecting the zymogen from autoprocessing.

**Figure 6.**
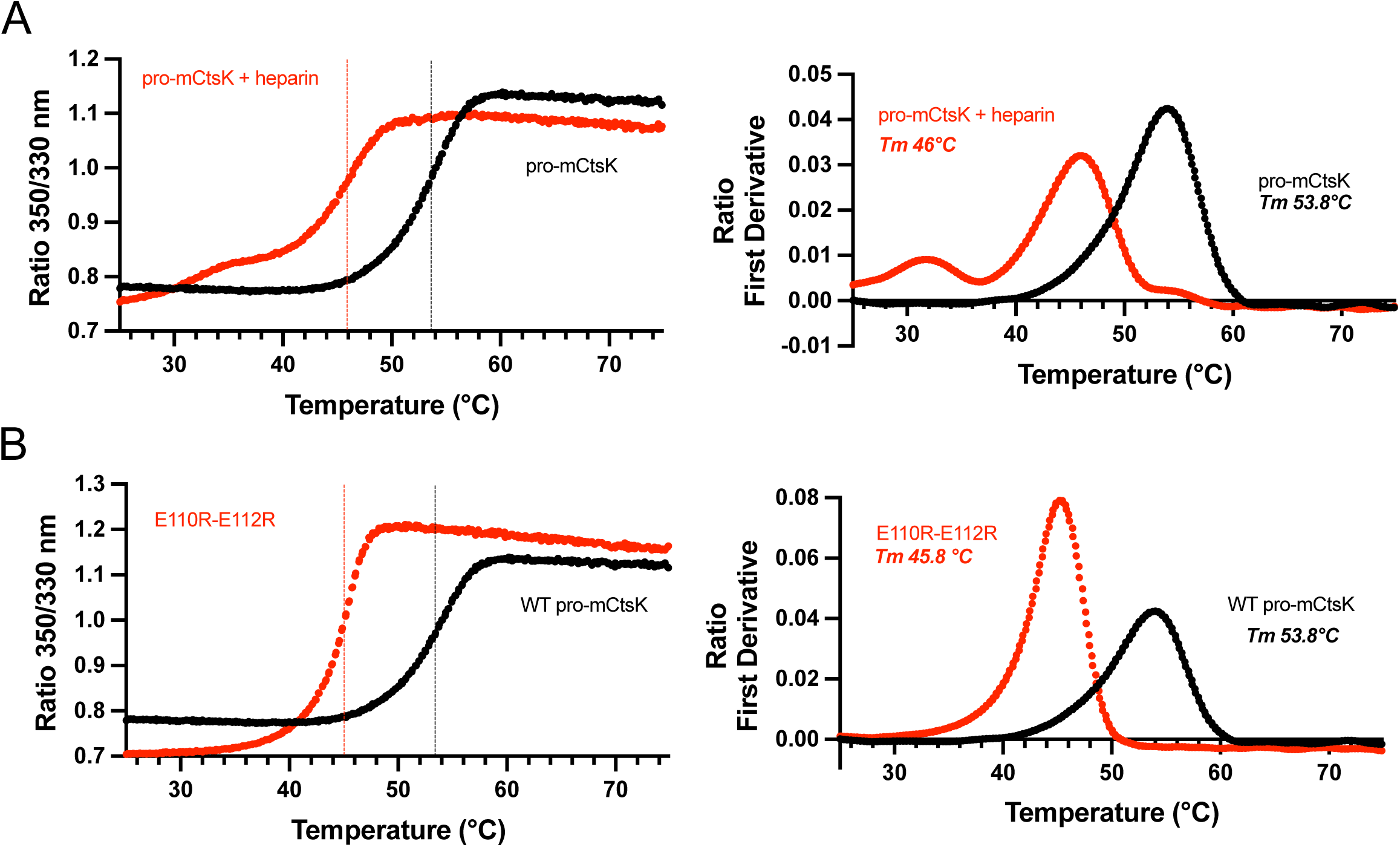
Disrupting the electrostatic interactions between the connection loop and the catalytic domain reduces the thermostability of pro-mCtsK. (A) NanoDSF thermograms of WT pro-mCtsK in the presence or absence of heparin at 1:1 molar ratio. Left, intrinsic fluorescence intensity ratio of tryptophan (350 nm/330 nm); right, first derivative of fluorescence ratio. (B) Thermograms of WT pro-mCtsK and E110R-E112R mutant. Data representative of three similar experiments.

### Mutating HS-binding residues away from the connection loop resulted in reduced effectiveness of HS-mediated autoprocessing

The fact that subtle changes in sulfation pattern and length of HS oligosaccharides resulted in clear differences in the rate of autoprocessing suggests that the systems is very sensitive to the affinity of HS–pro-mCtsK interaction (Fig. 2). Here we decided to further strengthen this conclusion by mutating the HS-binding residues of pro-mCtsK. To this end we mutated R222 and K328, which are also involved in HS-binding (Fig. 3A) but located farther away from the connection loop (Fig. 7A) (15). R222A-K328A double mutant resulted in a 75 mM reduction in the salt concentration required for elution from heparin Sepharose column, suggesting that these two residues are involved in HS–pro-mCtsK interaction (Table I). Unlike R225, R237, and R241, which are directly involved in interacting with the connection loop, we expect that mutating R222 and K328 has no direct effect on the autoprocessing of pro-mCtsK in the absence of HS. Indeed, R222A-K328A double mutant display similar rate of autoprocessing as WT pro-mCtsK in the absence of HS by 6 hrs (Fig. 7B, compared Fig. 1A). Interestingly, the enhancement effect of HS on the autoprocessing of R222A-K328A was reduced by more than 2-fold compared to WT (Fig. 5B), again suggesting the strength of HS–pro-mCtsK interaction determines the effectiveness of HS-mediated acceleration of autoprocessing.

**Figure 7.**
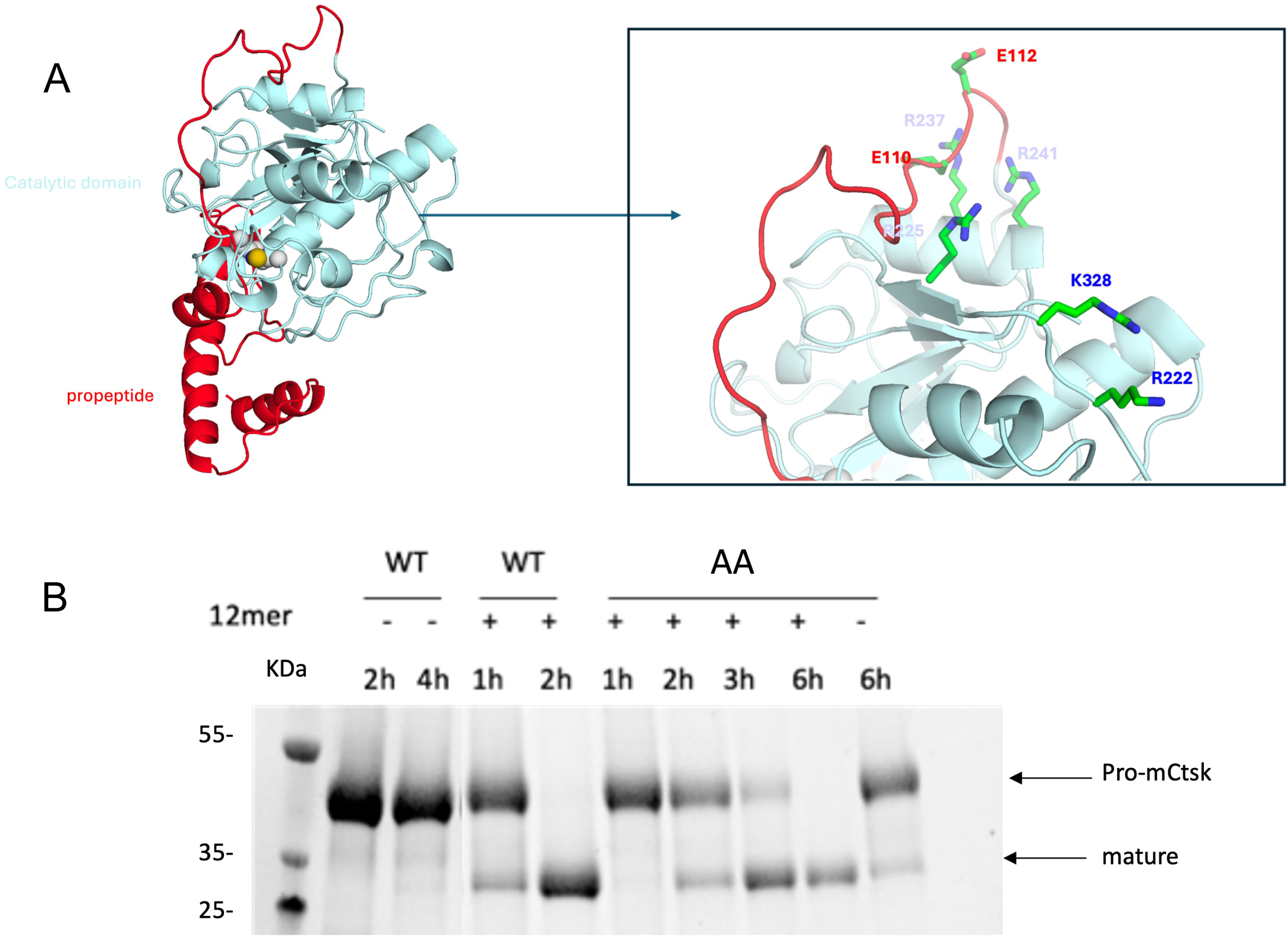
Weakening pro-mCtsK/HS interactions resulted in reduced responsiveness to HS-induced autoactivation. (A) Cartoon representation of the pro-hCtsk structure (PDB:1BY8) showing R222 and K328 are located far away from the connection loop containing E110 and E112. The propeptide is shown in red and the catalytic domain in cyan. B) Autoprocessing of WT and R222A-K328A (AA) mutant pro-mCtsK in the presence (+) or absence (-) of 12mer HS oligosaccharide at 22 C, pH 4.

### Co-localization of HS and CtsK in secretory lysosome of Osteoclasts

Previous studies suggest that secretory lysosomes contribute to both the storage and activation sites of CtsK, which eventually transport CtsK to the ruffled border for secretion into the resorption pits (16–18). Our results reveal a potential role of HS in promoting the autoactivation process of CtsK. Previously, we have shown that osteoclasts express high levels of CtsK and HS and they display extensive co-localization (15). To investigate whether HS participates in CtsK processing, we performed immunofluorescence staining of murine femur sections to visualize whether HS and CtsK colocalize in secretory lysosomes and ruffled border of osteoclasts. We co-stained bone sections with lysosomal marker LAMP2, which is known to be highly enriched in the secretory lysosomes and the ruffled border of osteoclasts (16). As shown in Figure 8B, we observed extensive co-localization of HS, CtsK and LAPM2 in osteoclasts. This finding suggest that HS is present in the same cellular compartment during the secretion of CtsK and likely contributes to CtsK maturation.

**Figure 8.**
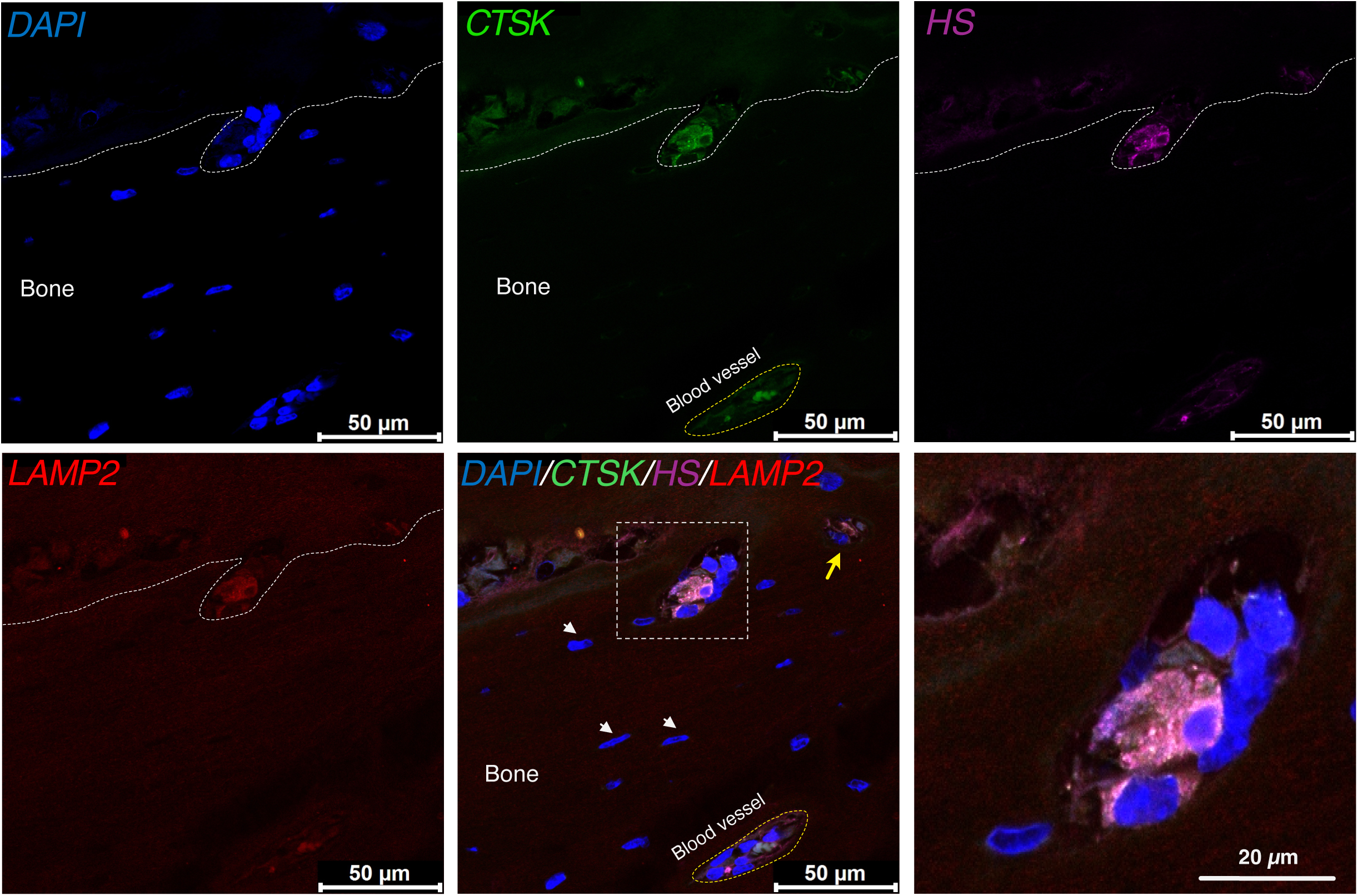
Co-localization of CtsK, LAMP2, and HS in osteoclasts. Mouse femur section was co-stained with anti-CtsK (green), HS20 (purple) and anti-LAMP2 (red) and visualized with anti-rabbit Alexa488, anti-human Alexa594, and anti-rat Alexa647 secondary antibodies, respectively. Area shown is the cortical bone. The approximate boundary of the bone is shown with white dashed line. A blood vessel within the bone is indicated with yellow dashed line. Red blood cells in the vessel shown strong green autofluorescence. A closeup view of the large osteoclast is shown in the right bottom panel. Another osteoclast (likely only the tip of the osteoclast) is indicated with a yellow arrow in the merged image. Nuclei of several osteocytes are indicated with white arrow head. Images representative of three separate experiments.

## DISCUSSION

Studies have shown that the rate of autoprocessing of several cysteine cathepsins, including cathepsin B, L and K, are greatly accelerated in the presence of GAGs(11, 13, 14). The molecular mechanisms that regulate this promotion effect of GAGs have been investigated in most detail in procathepsin B. Mutagenesis study of procathepsin B identified several basic residues in the propeptide, including His28, Lys39 and Arg40 (13), might be directly involved in binding to short HS oligosaccharide. They postulate that binding of short HS oligosaccharide to these residues induces a conformational change of the propeptide, turning it into a better substrate for intermolecular proteolytic activation. However, they also found that these residues are not involved in binding to C4S and regulating C4S-mediated acceleration of autoprocessing, suggesting full length GAGs might bind to a different site on procathepsin B and regulate autoprocessing through a different mechanism. Another study examined the mechanism by which GAG promote autoprocessing of procathepsin L. Through mutagenesis they identified two basic residues in the propeptide (K99 and K104) involved in binding to heparin and postulated that binding of heparin to the propeptide alters its conformation and facilitate proteolytic removal of the propeptide (14). However, in this study, the autoprocessing rate of the double mutant was not compared to the WT procathepsin L, making it unclear whether these two residues are truly involved in autoprocessing.

Compared to these earlier studies on the cathepsin B and L, our study revealed substantially more structural insights on the detailed mechanism by which GAGs promote autoprocessing of CtsK. Our study benefited from the recently solved co-crystal structure of mature CtsK and HS oligosaccharide, which clearly revealed the HS-binding site of CtsK, an information not available for cathepsin B and L. Overlay of the mature CtsK-HS co-crystal structure and the structure of proCtsK revealed a steric clash between the bound HS oligosaccharide and the connection loop of the propeptide (Fig. 3B). This observation suggests that stable HS-proCtsK interaction would almost certainly alter the conformation of the connection loop. Because the site of the steric clash is so close to the cutting site for removing the propeptide (between R114-A115), it is likely HS-binding renders the connection loop highly labile and becomes more susceptible for intermolecular proteolytic processing. Because residues R225, R237, and R241 (equivalent to residues R111, R123 and R127 when counting from the start of the mature enzyme ) make prominent contribution to proCtsK-HS interaction (15), we initially thought that mutating these residues might make proCtsK less sensitive to HS-mediated acceleration of autoprocessing. Unexpectedly, we found that R225A-R237A-R241A triple mutant display accelerated autoprocessing compared to WT proCtsK in the absence HS (Fig. 4B). A logical explanation for this observation is that these residues are directly involved in stabilizing the conformation of the connection loop to confer resistance to proteolysis. This led us to propose that the connection loop might be stabilized by the electrostatic interaction between E110 and E112, two conserved acidic residues in the connection loop, and R225, R237, and R241 (Fig. 4A). This model is supported by further mutagenesis study that introduces opposing charges to these residues (Fig. 4C), which resulted in even greater acceleration of autoprocessing, indicating greater destabilization of the connection loop compared to R225A-R237A-R241A mutant.

While our study provided the first structural insight of GAG-mediated acceleration of procathepsin autoprocessing, whether this same mechanism exists in other cathepsins remain to be tested experimentally. Our structural analysis of the crystal structure of procathepsin B and L revealed highly similar arrangement of acidic residues on the connection loop and basic residues on the same α-helix in the catalytic domain. It is possible that GAGs use a similar mechanism to accelerate autoprocessing of procathepsin B by binding to a similar binding site as proCtsK. The same mechanism might also in play in autoprocessing of procathepsin L. But procathepsin L is different from proCtsK in that several basic residues in its connection loop (which are not present in proCtsK) are directly involved in binding to GAGs (14). This difference suggests that the HS-binding site of procathepsin L is quite different from proCtsK, which in turn might lead to a somewhat different mechanism of destabilizing the connection loop.

To put these findings into physiological context, it is necessary to determine whether proCtsK and HS can direct interact with each other in the cell. Previous studies suggest that osteoclasts employ specialized lysosomes called secretory lysosomes for packaging and delivery of CtsK to the ruffled border for secretion (16–18). Immunostaining of CtsK revealed that it is highly enriched in secretory lysosomes and at the ruffled border (16). The observed extensive co-localization of HS, CtsK and lysosomal marker LAMP2 in mouse osteoclasts strongly suggest that HS and CtsK co-exists in secretory lysosomes and ruffled border (Fig. 8B). It is conceivable that when proCtsK is packaged into the secretory lysosome, it binds HS and become activated during the secretion process. Because the pH of lysosomes is usually maintained between 4.5 and 5, the rate of autoactivation of proCtsK would be very slow in the absence HS. Under this pH, the presence of HS might be a critical rate-limiting factor in determining the final yield of mature CtsK. Because we have shown that subtle changes in the sulfation level could have a profound effect of how fast the proCtsK is autoprocessed (Fig. 2), cellular regulation of HS biosynthesis in osteoclasts might also have an impact on how much mature CtsK is eventually produced. In addition, our finding that HS is highly enriched in the secretory lysosome suggests that HS might be an important component of osteoclast secretory machinery. It is possible that lysosomal HS in osteoclasts might play additional roles in osteoclast biology, which remains to be investigated.

## MATERIEALS AND METHODS

### Materials

Heparan sulfate (HS) oligosaccharides of defined lengths (10-, 12-, and 14-mer) and various sulfation patterns were obtained from Glycan Therapeutics (Raleigh, NC). Pharmaceutical-grade porcine heparin was purchased from Scientific Protein Laboratories. Chondroitin sulfate A (C4S, the major form in bone) was obtained from Millipore-Sigma. The fluorogenic cathepsin substrate Z-Leu-Arg-AMC was purchased from G-Biosciences. Unless otherwise stated, all chemicals were analytical grade, and all buffers were prepared in ultrapure water.

### Expression and purification of full-length murine pro-CtsK in mammalian cells

Complete open reading frame of full-length murine pro-CtsK was cloned into the pUNO1 mammalian expression vector (Invivogen) using AgeI and NheI restriction sites. Recombinant protein was expressed in 293-freestyle cells (Thermo Fisher Scientific) maintained in FreeStyle^TM^293 medium at 37 °C with 8% CO_2_ and shaking at 130rpm. Transient transfection was performed using FectoPRO (Polyplus) according to the manufacturer’s recommendations. Conditioned medium was harvested 5 days post-transfection, clarified by centrifugation (5000 rpm, 10 min) and 0.22um filtration, and loaded onto a HiTrap heparin-Sepharose column (Cytiva) equilibrated in 25mM HEPES, pH7.1 buffer. Bound protein was eluted using a 0-2M NaCl gradient. After purification, Pro-mCtsK was >99% pure as judged by silver staining.

### Site-directed Mutagenesis

Pro-mCtsK mutants were prepared using a previously published method (19). Mutations were confirmed by Sanger sequencing. Expression and purification of the mutants was performed the same way as described for *wild-type* (WT) Pro-mCtsK.

### Autoprocessing of Pro-mCtsK

Autoprocessing assays were performed to assess the conversion of the proenzyme to its mature form. Purified pro-mCtsK (typically 1 mg/ml unless indicated otherwise) was incubated in 100 mM sodium acetate, 8 mM EDTA, pH4, at room temperature (RT). For glycosaminoglycan-induced activation, reactions contained a 1:1 molar ratio of pro-mCtsK to glycosaminoglycan (heparin, C4S, or HS oligosaccharides of various lengths and sulfation levels). For pH-dependence studies, identical reactions were performed across buffers adjusted to pH 4-9 without adding GAGs. The reactions were terminated at the indicated time points by adding 10 µM E-64, a covalent cysteine protease inhibitor, followed by immediate boiling for 5min in 4× LDS loading buffer. Samples were separated on 4-20% Bis-Tris SDS-PAGE gels (GenScript) and visualized with Coomassie Blue.

### Peptidase activity after processing

CtsK enzymatic activity was measured using the fluorogenic substrate Z-Leu-Arg-AMC in a fluorescence microtiter plate. After Pro-mCtsK was incubated in the autoactivation buffer described above, 1ul of each reaction was diluted into 50ul of assay buffer (100 mM sodium acetate, 2.5mM EDTA, pH5.5). The diluted enzyme was combined with an equal volume of substrate solution (100 µg/ml). The fluorescence signal (excitation: 370nm, emission: 450nm) was monitored in black 94-well plates using a SpectraMax plate reader (Molecular Devices) for 30min with 30 seconds intervals. Activity was calculated from the linear portion of the reaction progression curve.

### Nano Differential Scanning Fluorimetry

NanoDSF was performed using Prometheus Panta (NanoTemper Technologies). 20 µl of purified WT pro-mCtsK and E110R-E112R mutant, both at 0.5 mg/ml (in 25 mM HEPES, 150mM NaCl, pH7.2), were mixed with 20 µl of 100mM sodium acetate, 8mM EDTA, pH3.9 to bring the final pH to 4 and final concentration to 250 µg/ml. The mixture was immediately loaded in nanoDSF standard capillaries and exposed at thermal stress from 25 °C to 80 °C by thermal ramping rate of 1 °C/min. For selected WT pro-mCtsK samples, heparin was added to final concentration of 60 µg/ml (1:1 molar ratio to pro-mCtsK) before loading into the capillaries. Fluorescence emission from tryptophan after UV excitation at 280 nm was collected at 330 nm and 350 nm. Fluorescence intensity ratio (350/330 nm) and Ratio First Derivative were calculated by ThermControl software.

### Immunohistochemistry

Femurs from 10-week-old C57BL/6 mice were collected, fixed in 10% neutral buffered formalin for 48 hours, and decalcified in 10% EDTA for 2 weeks at RT. Samples were embedded in paraffin and sectioned at 5um. Following deparaffinization and citric acid-based antigen retrieval (pH7), sectioned were blocked and incubated overnight at 4 °C with the following primary antibodies: 0.2 µg/ml rabbit anti-mCtsK polyclonal antibody (described previously) (15), 1ug/ml human anti-HS mAb (HS20, from Bio X cell) (20), and 1ug/ml Rat anti-mouse Lamp2 (GL2A7, Developmental Studies Hybridoma Bank). For negative controls, sections were treated with 5 mu/ml of heparin lyase III for 1hr at RT and incubated with species matched IgG controls. For immunofluorescence staining, slides were treated with anti-rabbit IgG Alexa Fluor-488 to visualize CtsK, anti-human IgG Alexa Fluor-594 to visualize HS, and anti-rat IgG Alexa Fluor-647 to visualize secretory lysosome. Nuclei were counterstained with DAPI. The images were collected with a Leica confocal microscopy (Stellaris 5) using a 63× oil objective. The images were processed using Leica Application Suite X software (Leica).

### Statistical analysis

All data are expressed as means ± SDs. Statistical significance was assessed of Variance (ANOVA) using GraphPad Prism software (GraphPad Software Inc.). P value<0.05 was considered statistically significant.

## ACKNOWLEDGMENTS

This work is supported by National Institutes of Health grant R01DE031273 (to DX, JL) and R01AR078212 (to DX).

## Reference

1. A. G. Costa, N. E. Cusano, B. C. Silva, S. Cremers, J. P. Bilezikian, Cathepsin K: its skeletal actions and role as a therapeutic target in osteoporosis. Nat Rev Rheumatol 7, 447–456 (2011).

2. P. Garnero et al., The collagenolytic activity of cathepsin K is unique among mammalian proteinases. J Biol Chem 273, 32347–32352 (1998).

3. M. H. Polymeropoulos et al., The gene for pycnodysostosis maps to human chromosome 1cen-q21. Nat Genet 10, 238–239 (1995).

4. W. S. Hou et al., Characterization of novel cathepsin K mutations in the pro and mature polypeptide regions causing pycnodysostosis. J Clin Invest 103, 731–738 (1999).

5. B. D. Gelb, J. G. Edelson, R. J. Desnick, Linkage of pycnodysostosis to chromosome 1q21 by homozygosity mapping. Nat Genet 10, 235–237 (1995).

6. Y. Xue et al., Clinical and animal research findings in pycnodysostosis and gene mutations of cathepsin K from 1996 to 2011. Orphanet J Rare Dis 6, 20 (2011).

7. S. Verma, R. Dixit, K. C. Pandey, Cysteine Proteases: Modes of Activation and Future Prospects as Pharmacological Targets. Front Pharmacol 7, 107 (2016).

8. M. S. McQueney et al., Autocatalytic activation of human cathepsin K. J Biol Chem 272, 13955–13960 (1997).

9. L. Zhang, Glycosaminoglycan (GAG) biosynthesis and GAG-binding proteins. Prog Mol Biol Transl Sci 93, 1–17 (2010).

10. M. Novinec, B. Lenarčič, B. Turk, Cysteine cathepsin activity regulation by glycosaminoglycans. Biomed Res Int 2014, 309718 (2014).

11. P. A. Lemaire et al., Chondroitin sulfate promotes activation of cathepsin K. J Biol Chem 289, 21562–21572 (2014).

12. K. K. Bojarski, A. S. Karczyńska, S. A. Samsonov, Role of Glycosaminoglycans in Procathepsin B Maturation: Molecular Mechanism Elucidated by a Computational Study. J Chem Inf Model 60, 2247–2256 (2020).

13. D. Caglic, J. R. Pungercar, G. Pejler, V. Turk, B. Turk, Glycosaminoglycans facilitate procathepsin B activation through disruption of propeptide-mature enzyme interactions. J Biol Chem 282, 33076–33085 (2007).

14. M. Fairhead, S. M. Kelly, C. F. van der Walle, A heparin binding motif on the pro-domain of human procathepsin L mediates zymogen destabilization and activation. Biochem Biophys Res Commun 366, 862–867 (2008).

15. X. Zhang et al., Heparan sulfate selectively inhibits the collagenase activity of cathepsin K. Matrix Biol 129, 15–28 (2024).

16. E. van Meel et al., Disruption of the Man-6-P targeting pathway in mice impairs osteoclast secretory lysosome biogenesis. Traffic 12, 912–924 (2011).

17. S. Ohmae et al., Actin-binding protein coronin 1A controls osteoclastic bone resorption by regulating lysosomal secretion of cathepsin K. Sci Rep 7, 41710 (2017).

18. P. Y. Ng, A. Brigitte Patricia Ribet, N. J. Pavlos, Membrane trafficking in osteoclasts and implications for osteoporosis. Biochem Soc Trans 47, 639–650 (2019).

19. L. Zheng, U. Baumann, J. L. Reymond, An efficient one-step site-directed and site-saturation mutagenesis protocol. Nucleic Acids Res 32, e115 (2004).

20. W. Gao, Y. Xu, J. Liu, M. Ho, Epitope mapping by a Wnt-blocking antibody: evidence of the Wnt binding domain in heparan sulfate. Sci Rep 6, 26245 (2016).

